## Supplementary material for "Genomic islands targeting *dusA* in *Vibrio* species are distantly related to *Salmonella* Genomic Island 1 and mobilizable by IncC conjugative plasmids": S2 Table

### S2 Table. Oligonucleotides used in this study.

| **Primer name** | **Nucleotide sequence (5' to 3')*^a^*** |
| --- | --- |
| dusAigEcoRIF | NNNNGAATTCACAAGTTATCGCTCTATCACTG |
| dusAigEcoRIR | NNNNGAATTCCCTTTGATGTGGGGCATG |
| oRD1 | CAGCCGACGTTGAGGTTAA |
| oRD2 | CAGCCGACATTCAGGTTG |
| oRD3 | AAGCGAATGATGCCTTTACTG |
| oRD4 | GTGTGTGTAGCTTCAGGTG |
| oRD5 | GTCGAACCTGGATTGTTTATCATTG |
| oRD6 | TCACCATCACTGTTGGACTT |
| dusAscarNoFRTf | CAGCAGTCCTATCATGCCCCACATCAAAGGGAATTCagagcgcttttgaagctca |
| dusAscarNoFRTr | GATCATAATCAATATCGATGGAGAAAAGCAATGACAtcggaataggaacttcaaga |
| DelprtNf | ATCGTTGGAAATTGTTGAGAATGATTGAGGATAGCTgtgtaggctggagctgcttcg |
| DelprtNr | GGGATGGGATAATATTTGGCATTCAGACCCAGGTAGTTAattccggggatccgtcgacc |
| FwDeltaDapA-MG1655 | ATGTTCACGGGAAGTATTGTCGCGATTGTTACTCCGGTGTAGGCTGGAGCTGCTTCG |
| RvDeltaDapA-MG1655 | TTACAGCAAACCGGCATGCTTAAGCGCCGCTCTGACCATATGAATATCCTCCTTA |
| lacZW-B | GCGAAATACGGGCAGACATGGCCTGCCCGGTTATTACATATGAATATCCTCCTTA |
| lacZW-F | TTGTGAGCGGATAACAATTTCACACAGGAAACAGCTGTGTAGGCTGGAGCTGCTTCG |
| oDF15 | AGCGTTGCACCAATGCTCGACTGGACGGACAGACATCTGGCCGTCGTTTTACAACGTCG |
| oDF16 | AGGGCGTGGTGAATTTGACTACTTTTTGGTGAAAAGGCAGCATTACACGTCTTGAG |
| oFD26f | gcagaacgggcattcgacacaagttcgctgattaacgtgtaggctggagctgcttc |
| oFD26r | CCAGGTCTTTGGCCGCAAAAATGAGGATGAGTAGTCcatatgaatatcctcctta |
| oFD1r | NNNNCTCGAGCACATGATTTCCGGAAATAAAAGC |
| oFD1f | NNCTGCAGTTAATCAGCGAACTTGTGTCGAAT |
| oFD3r | NNNNCTCGAGAGACAAATACTCCCGACTTGATCC |
| oFD4r | NNNNCTCGAGAGCTATCCTCAATCATTCTCAACA |
| oFD4f | NNCTGCAGTAAAAACATTTGAGAGGTCATTCGG |
| oFD5r | NNNNCTCGAGAGACACCTCCAAAAAGTTGAAGG |
| oFD5f | NNCTGCAGGCAGCTTATAGCATGAATCTGTAC |
| oFD6r | NNNNCTCGAGGAAGCGAATGATGCCTTTACTGG |
| oFD6f | NNCTGCAGCGCTGAATCTACGACTTAATGACA |
| prtNEcoRIf | NNNGAATTCAAGGAGGAATAATAAATGAATACCGCATTTCTTCTG |
| prtNHindIIIrev | NNNaagcttCAGGTAGTTACAGTTCTCTC |
| oFD38r | NNNaagcttTGAAACTGCCCATTTTGGGAA |
| oFD44f | NNNgagctcCAATACCGCCAGGTCTTTGG |
| oVB10 | CAGGTGGCACTTTTCGGGGTCAGTCTTTCCCAACACTCATCCCCTTCTG |
| oVB11 | GAGTGTTGGGAAAGACTGACCCCGAAAAGTGCCACCTGCATCGATG |
| oVB12 | NNNGAGCTCCGGGCGAGAAGTAGCGTTGA |
| oVB13 | NNNGAGCTCGATAGCTAGACTGGGCGGT |
| attBdusAqPCRfwd | CCTGAATAGTGATGCTGAATAAC |
| attBdusAqPCRrev | CCATTTCGGTATACAGCAAC |
| higAqPCRfwd | CTTCCTGCTCAAAGACTCTATG |
| higAqPCRrev | CTGGTGACCGAGTTTCTG |
| qFwpVCR | AAGAGAACCAAAGACAAAGACC |
| qRvpVCR | CACCTTCACCGTGAAATGC |
| qdnaBFw | ACGATTTTTACACCCGCCCAC |
| qdnaBRv | ATCATCTCACGGACAACGGCAC |
| qhicBFw | GCTTATCCCTTTACCTTCGCC |
| qhicBRv | TAACTCTTTGCCAAGCGCC |
| qthdFFw | GATAATGACACTATCGTAGCCC |
| qthdFRv | GCAGTTCCAGCACATCTTC |

*^a^* Cloning sites are underlined
