## Supplementary figures and images for "Genomic islands targeting *dusA* in *Vibrio* species are distantly related to *Salmonella* Genomic Island 1 and mobilizable by IncC conjugative plasmids"

### S1 Fig

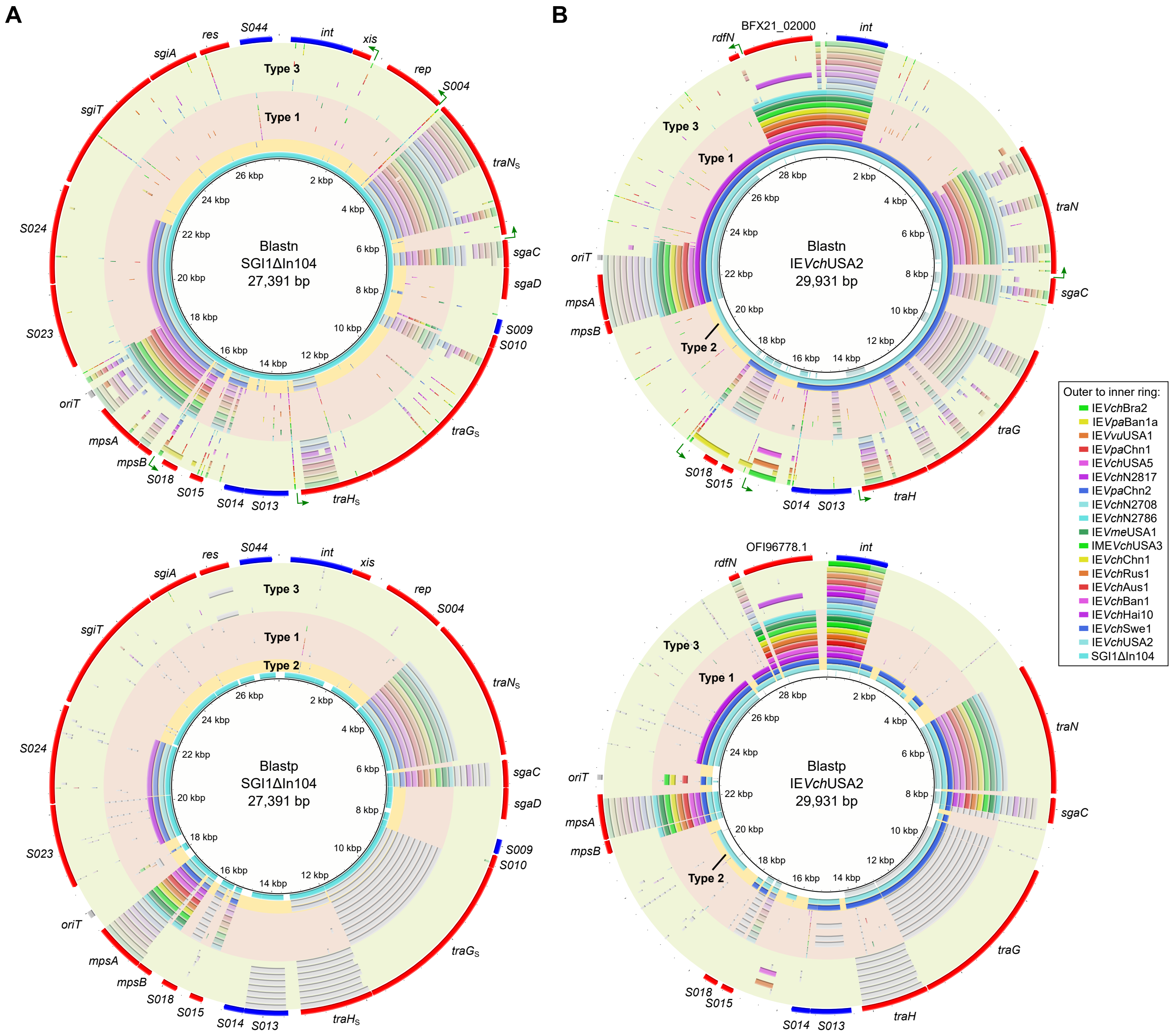

### S2 Fig

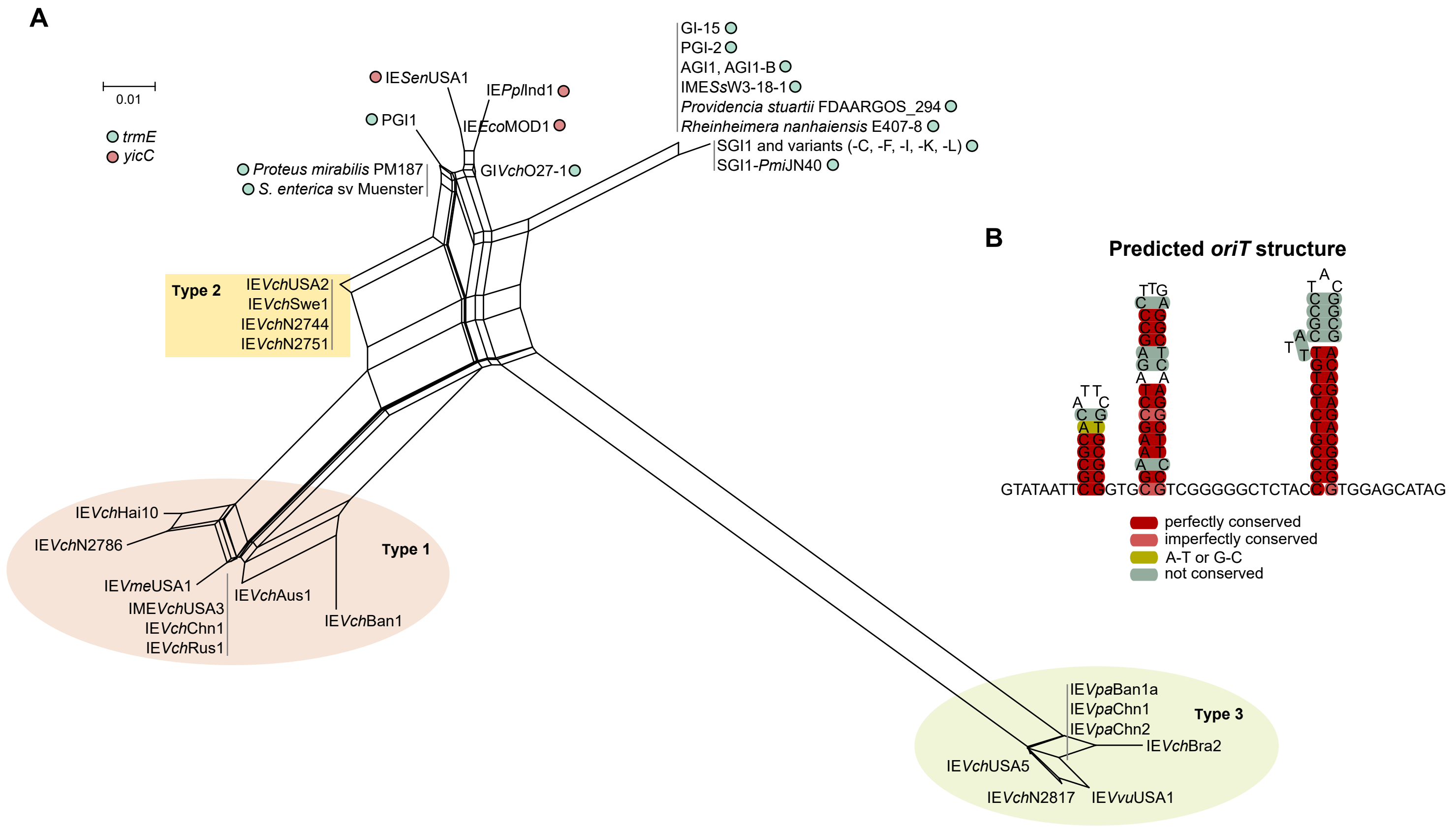

### S3 Fig

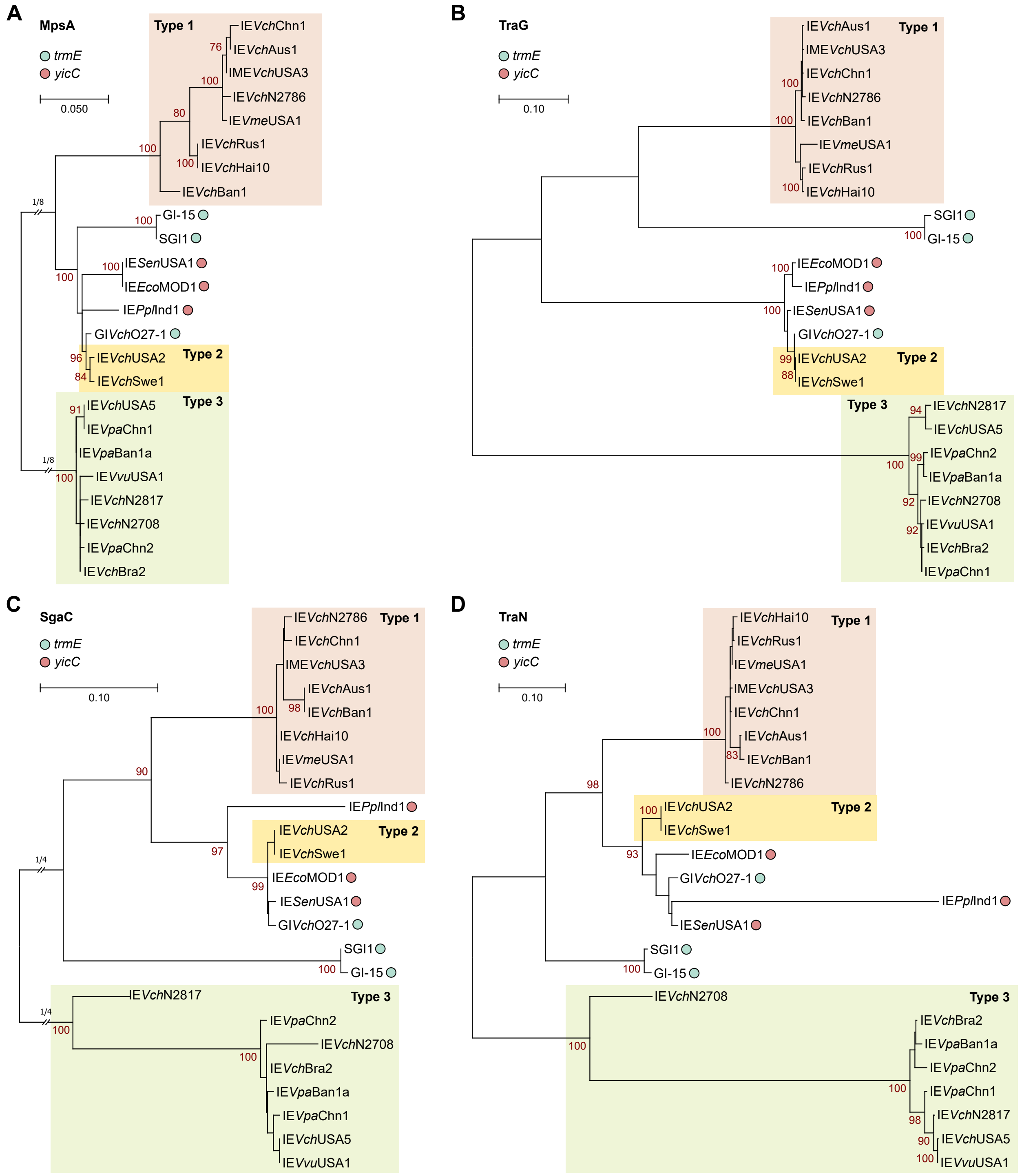

### S5 Fig

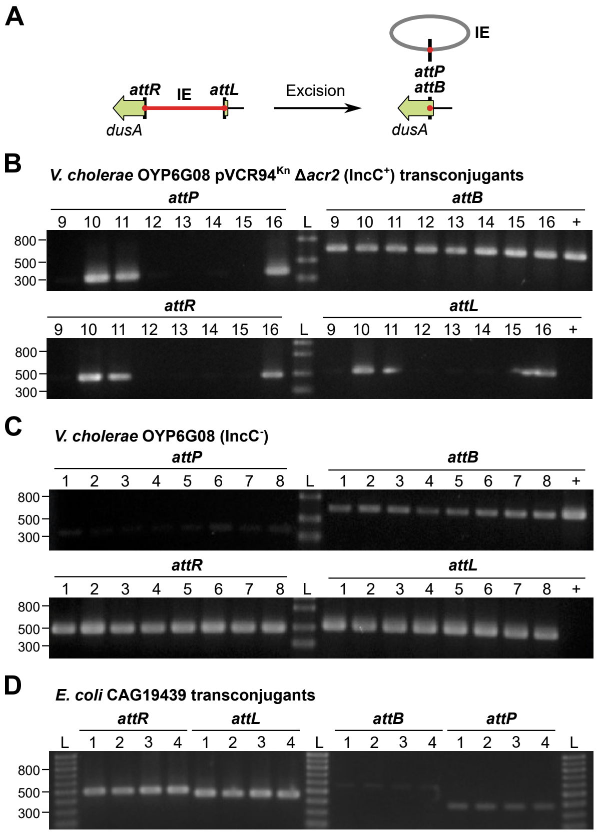
