## Supplementary material for "Genomic islands targeting *dusA* in *Vibrio* species are distantly related to *Salmonella* Genomic Island 1 and mobilizable by IncC conjugative plasmids": S4 Fig

**A**

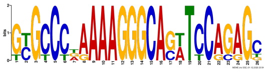

| | | ***p*-value** |
|---|---|---|
| SGI1 *xis* | TTC**GCGCCCTAAAAGGGCAGATCCAGAGC**CGAGGATTGGTAGCTGTCTCTCATAAACTAGGTCTATCAATAACTGGTGTGTAACGGGTAGGCTG**ATG** | 4.2E-10 |
| GI-15 *xis* | TTC**GCGCCCTAAAAGGGCAGATCCAGAGC**CGAGGATCTGAAACCTGTGTTGGTAGACTAAACCAG(22)AGATATGACTATGCAAGAGACAGAG**TTG** | 4.2E-10 |
| GI*Vch*O27-1 *xis* | GAG**TTGTCCAAAAAGGGCACTTTCAGAGC**GGAGGTTTCTCAACGGGCACCCATAAAATTGCAGTC(6)AGGGTTGCCAATGAAGTAGGTATGCG**ATG** | 6.8E-08 |
| IE*Eco*MOD1 *rdfM* | TTT**GCGCCCGAAAAGGGCACTTCCAGAGG**TGAGGTTTTTAGACGCCACCCAGCAAAATTGCCCCTGTGATCAAATTCATCAGGGGAACCT**ATG** | 6.8E-12 |
| IME*Vch*USA3 *rdfN* | GTG**GTGCCCGAAAAGGGCAGTTCCAGAGC**CGATGATTTGCAGTGTTGAACCTTAGGATCGTTGGAAATTGTTGAGAATGATTGAGGATAGCT**ATG** | 4.1E-11 |
| IE*Vch*USA5 *rdfN* | TAA**GTGCCTAGAAAGGGCAGATCCGCGGT**ATGTCTCGCATGTTGAAATAAACGAAAATGATGGCGTTTAGTTGAGGA**ATG** | 6.5E-08 |

**B**

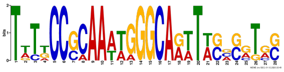

| | | ***p*-value** |
|---|---|---|
| SGI1 *traN*$_S$ | CG**TATGCCGCAAAAGGGCAAATAGCGATGC**TAATTTTTATGAGAGAGCGACATAGCATTATCCAAACTAAAAAGCTGGAGAAATGCT**ATG** | 6.4E-06 |
| GI-15 *traN* | CG**TATGCCGCAAAAGGGCAAATAGCGATGC**TAATTTTTATGAGAGAGCGACATAGCATTATCCAAACTAAAAAGCTGGAGAAATGCT**ATG** | 6.4E-06 |
| GI*Vch*O27-1 *traN* | GA**TTTTCCGCAATTGGGCAGTTTGGCGTAG**GAGGTTTATTACTTCAGCCGAATAGCATAGCTCTCAAGATTTTATTGAGAGCTATGCT**ATG** | 5.4E-06 |
| IE*Eco*MOD1 *traN* | GA**TTTTCCGCAATTGGGCAGTTTCGCGTAG**GAGGTTTATTACTTCAGCCGAATAGCATGGTTCTCAAGATTTTATTGAGAGCCATGCT**ATG** | 3.7E-06 |
| IME*Vch*USA3 *traN* | TG**TTTTCCCAAAATGGGCAGTTTCACCGCG**TAGGTTTTGGTGGGTGAGCTTTTAACATGATTTTCTAACGCTTTTATTTCCGGAAATCATGTG**ATG** | 2.3E-08 |
| IE*Vch*USA5 *traN* | AA**TGCACCCCAAATAGGCAATTACAGGTGC**GTAAGAATTTTAAATACTGAAATAGCATGAACCATTGAAAATGGAGGTATCATCT**ATG** | 2.8E-08 |

**C**

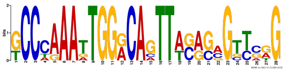

| | | ***p*-value** |
|---|---|---|
| SGI1 *traHG*$_S$ | GGACT**GCCCAAATTGGACAGTTTGGGAGTTCGG**TTTGTACTCATTTGCTCGGTAGGCTTTCGGGTGACACGAAACCTATTGGAGCAACAGT**ATG** | 8.9E-09 |
| GI-15 *traHG* | GGACT**GCCCAAATTGGACAGTTTGGCAGTTCGG**TTTGTACTCATTTGCTCGGTAGGCTTTCGGGTGACACGAAACCTATTGGAGCAACAGT**ATG** | 7.9E-09 |
| GI*Vch*O27-1 *traHG* | AGGAT**GCCCAAAATGGACACTTACAGCGTTCGG**TTTGCCGATTATCGGTGGTTACTCTGGAGCCAACGACTATCCGCAAATTGAGGTTGAA**ATG** | 2.5E-10 |
| IE*Eco*MOD1 *traG* | TAAAC**GCCAAAATTGGGCAGTTACAGAGTCGTG**GTAAAGAGGATTGAGCGAATAGGCTATCTCCAAATTTCGGGAGAATAAGCCT**ATG** | 5.6E-11 |
| IME*Vch*USA3 *traG* | ATTAT**TCCTGAATTGGGCAGTTACACGGGCGAG**GTTTGAAGCCAAATTGCGATAGGATGAACGGATCAAGTCGGGAGTATTTGTCT**ATG** | 8.2E-07 |
| IE*Vch*USA5 *traHG* | ATTGC**TCCCAAAGTGGGCAATTAGAGCGCTAAG**GAATAGCGCTAATGATTGAGAAAATAGAAACTCAAATTCGGATGGAGAGTTTCTATT**ATG** | 6.0E-07 |

**D**

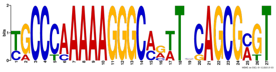

| | | ***p*-value** |
|---|---|---|
| SGI1 *S018* | AATG**TGCCCAAAAAGGGCCAATACAGCGCGT**GTCGATTCAGAAATGAGCTTGTTAAATTTGAATC(110)AATTCCAACTAATTGGAGGTGTTT**ATG** | 2.8E-12 |
| GI-15 *S018* | ATTG**TGCCCAAAAAGGGCCAATACAGCGCGT**GTCGATTCAGAACTGAGCTGTTAAATTTGAATTA(109)AATTCCATCTAATTGGAGGTGTTT**ATG** | 4.7E-12 |
| GI*Vch*O27-1 *S018* | TATG**CGCCCAAAAAGGGCAGTTCCAGCGAGT**ATCCCCTGACGCCTTGGCTGTTACAGTGTTCACA(101)CCTCAAAGAAATTTGAGGTGTTGT**ATG** | 3.2E-13 |
| IE*Eco*MOD1 *S018* | ACGA**TGCCCAAAAAGGGCAGTTCCAGCGAGT**ATCCTCTGACGTATTGCTTGTTAAAGTGTTCTCA(101)CCTCAAAGAAATTTGAGGTGTTGT**ATG** | 1.9E-08 |
| IME*Vch*USA3 *S018* | CGTT**TACCCCAAAAGGGCACTTTCAGCGCGT**TCCATCACATCCCATCTTTTATAAACTATCCACA(101)CCCTTCAACTTTTTGGAGGTGTCT**ATG** | 1.0E-07 |
| IE*Vch*USA5 *S018* | AGTG**CGCCTAAAAAGGGCAGTTGGAGCGATT**TGTCTAAGTAGAACAGCCGGTTATCTTGGAAGTT(104)CCCTCAAAAATATGAGGTGAAGCT**ATG** | 1.4E-06 |
